# Neurodevelopmental Inequality arising from Early Childhood Stunting: Evidences from Brain Connectivity

**DOI:** 10.64898/2026.06.08.730782

**Authors:** Samuel Berkins, Suman Saha, Anitha Jasper, Roshan Livingstone, Sushil John, Venkata R Mohan, Beena Koshy, Arpan Banerjee

## Abstract

Early childhood stunting (ECS) affects millions of children globally and conjectured to result in suboptimal brain and cognitive development later in life. Charting out the trajectory of brain network development and most importantly how the compensation of function can be achieved gives windows of intervention to clinicians, educators and policy makers. In this study, advanced network neuroscience tools, graph theoretical methods applied on diffusion weighted MR-imaging (DWI) revealed the effects of ECS on white matter (WM) organization in a community-based birth cohort of 170 children (mean age = 9.18 years, SD = 0.28) from Vellore, India. Based on stunting status at ages two, five, and nine years, children were categorised into four groups: always stunted (AS; ***n***=21), stunted until five with catch-up at nine (S5C9; ***n***=31), stunted until two with catch-up at five (S2C5; ***n***=28), and never stunted or typically developing (TD; ***n***=90). The catch-up groups showed strikingly similar anthropometric measures compared to TD. However, all stunted groups (AS, S5C9, S2C5) showed significantly lower performance and verbal IQ scores compared to TD children. DWI data revealed AS children exhibited shorter tract lengths across select cortico-cortical connections, an increased number and strength of short- and medium-range connections, and a corresponding reduction in long-range connections and strength. Network analyses further revealed that AS and S5C9 groups displayed higher local clustering and local efficiency relative to TD, reflecting greater local segregation of brain networks. Hub-like modules was broadly conserved across groups, although both catch-up groups (S2C5 and S5C9) showed additional hubs, suggesting compensatory network reorganization via a more modular architecture. Together, these findings provide novel evidence that ECS is linked to altered structural reorganization of brain networks and reduced cognitive performance in later childhood. While persistent stunting (AS) is associated with the most pronounced alterations, partial catch-up (S2C5 and S5C9) is accompanied by compensatory adaptations, such as increased short-range connectivity and recruitment of additional hubs. These results underscore the critical importance of early nutritional interventions to support optimal brain network development.

## 1 Introduction

Early childhood stunting (ECS) remains a major global public health concern, affecting over 150 million children worldwide (World Health Organization, 2023). Defined as height-for-age more than two standard deviations below World Health Organization growth norms, ECS involves chronic disruption to early physical growth trajectories (World Health Organization, 2006). Beyond impaired somatic growth, ECS is consistently associated with impaired neurocognitive behaviours and suboptimal linguistic performances [1–5]. Despite extensive documentation of these functional outcomes, the synaptic pathways which impact early developmental milestones are poorly understood. This is also a major lacunae in development of clinically informative neuroimaging markers of ECS and its associated atypical developmental trajectories. Hence, bridging the knowledge gap by charting developmental trajectory along the axis of brain structural complexity will be of immense value to different stakeholders - basic scientists, developmental paediatricians and public health researchers.

Evidence linking early childhood malnutrition and stunting to the atypical brain development is sparse and methodologically heterogeneous [6]. Also, neuroimaging studies using advanced structural MRI are relatively rare and mainly focusing on macrostructural brain morphology rather than neuronal pathways, quantified by structural connectivity (SC) from white matter fiber counts between any pair of brain regions. For instance, reduced total brain volume has been reported in adolescence with a history of childhood malnutrition [7]. More recent studies have yielded mixed findings, with some reporting minimal or inconsistent morphological differences after recovery, often in relatively small or partially imaged samples [8]. Another work on the same cohort has refined this picture, demonstrating graded reductions in total brain volume, subcortical structures, cerebellar white matter, and posterior corpus callosum thickness across children with normal growth, persistent stunting, and catch-up growth, alongside regionally specific reductions in occipital and basal ganglia volumes [9]. Critically, despite emerging evidence for morphometric alterations associated with early growth disruption, no studies to date have directly characterized how ECS relates to white matter organization, the brain-wide structural connectivity, or the spatial architecture of brain networks, limiting mechanistic understanding of how early physical growth failure becomes embedded within distributed neural systems.

Addressing the research gap requires understanding normative white matter development and its sensitivity to early-life conditions that relies on prolonged axonal growth, progressive myelination, and activity-dependent refinement extending from infancy through adolescence [10, 11]. Beyond microstructural maturation, typical development involves systematic reorganization of brain-wide neural connectivity, with a shift from predominantly local connections toward greater long-range integration [12, 13]. This process supports the emergence of highly connected hub regions that facilitate efficient communication between distributed cortical and subcortical systems [14]. These hubs preferentially interconnect through rich-club architecture, forming a backbone for global integration while balancing metabolic and wiring costs [15]. To quantify these developmental changes in large-scale network organisation, the graph-theoretical metrics provide a principled framework. Structural brain networks typically exhibit small-world topology, characterised by high local clustering alongside short communication paths that support efficient information transfer [16–18]. Across childhood and adolescence, increasing integration, stabilisation of hub architecture, and refinement of long-range connections accompany cognitive maturation [14, 19, 20]. Disruptions to these network-level properties, including alterations in segregation, integration, or hub structure, have been implicated across neurodevelopmental and psychiatric conditions, highlighting network organisation as a sensitive marker of atypical development [21, 22]. Yet whether ECS is associated with similar alterations in white matter network architecture, and whether subsequent catch-up growth reflects normative recovery or compensatory reorganisation, remains unknown.

In the present study, diffusion MRI and graph-theoretical metrics was used to investigate how early childhood stunting and subsequent catch-up growth are associated with the organization of white matter structural networks in late childhood. Tract-level properties, the spatial distribution of short- and long-range connections, global and nodal network topology, and hub architecture were examined.

## 2 Methods

### 2.1 Participants

The present study is a subset of the multinational longitudinal prospective cohort “Etiology, Risk Factors and Interactions of Enteric Infections and Malnutrition and the Consequences for Child Health and Development” Network [23], investigating the risk and protective factors for brain development in children in India. The original birth cohort recruited 251 newborns from eight heavily populated urban slum residences in Vellore, South India, between March 2010 and February 2012. The cohort was subsequently followed up at different time points and MRI was acquired at nine years of age. Further details of the original Indian study population, recruitment, inclusion and exclusion criteria can be found in the Refs [3, 24, 25]. Families with existing migration plans, multiple pregnancies, another child already enrolled in the study and medical comorbidities were excluded. Out of the 251 children, only 205 children were available for the nine-year follow-up. Among these 205 children, the parents of 191 children agreed to be available for MRI brain scans. Of these, six children did not cooperate for the MRI scan. The remaining 185 children underwent MRI brain scans between 2020 and 2021. Of which, one child had severe motion artefacts, four children did not fall into any group, eight MRI datasets were incomplete, and two children had low intelligence quotient. Following these exclusions, 170 children with a mean age of 9.18 years (ranging from 8.1 to 10.8 years; M/F: 78/92) were included in the final analysis (Table 1). Of which, 90 children were never stunted (NS), 28 children were stunted till two with catch-up at five (S2C5), 31 children were stunted till five with catch-up at nine (S5C9) and 21 children who were always stunted (AS). All children were born at or after 37 gestational weeks (Figure. 1a-b). The Institutional Review Board and Ethics Committee of Christian Medical College, Vellore, India approved the initial birth cohort and all further follow-ups. Written informed parental consent was obtained at all time points, along with the child’s assent at nine years.

**Table 1.** Demographic and anthropometric characteristics of the study sample.

|  | NS (N = 90) | S2C5 (N = 28) | S5C9 (N = 31) | AS (N = 21) | p-value<sup>2</sup> |
| --- | --- | --- | --- | --- | --- |
| **Sex, n (%)** |  |  |  |  |  |
| Female | 57 (63) | 13 (46) | 13 (42) | 9 (43) | 0.083 |
| Male | 33 (37) | 15 (54) | 18 (58) | 12 (57) |  |
| **Characteristics at 2-year follow-up** |  |  |  |  |  |
| Height (cm) | 83.0 (81.7–84.4) | 80.1 (78.9–81.0) | 78.5 (77.5–80.0) | 76.4 (75.2–77.1) | < 0.001 |
| Weight (kg) | 10.4 (9.9–11.1) | 9.56 (9.8–10.2) | 9.2 (8.7–9.6) | 8.9 (8.1–9.6) | < 0.001 |
| **Characteristics at 5-year follow-up** |  |  |  |  |  |
| Height (cm) | 105.0 (102.8–107.0) | 102.2 (101.2–104.0) | 98.9 (97.9–99.3) | 95.2 (93.7–97.4) | < 0.001 |
| Weight (kg) | 15.6 (14.6–16.8) | 14.7 (14.2–15.6) | 13.7 (12.7–14.5) | 12.8 (12.1–13.2) | < 0.001 |
| **Characteristics at 9-year follow-up** |  |  |  |  |  |
| Age (years) | 9.2 (8.1, 9.8) | 9.2 (9.0, 10.8) | 9.11 (8.9, 9.6) | 9.2 (9.0, 10.3) | 0.55 |
| Height (cm) | 131 (129–135) | 129 (126–131) | 123 (122–125) | 118 (117–120) | < 0.001 |
| Weight (kg) | 26 (23–30) | 24 (23–28) | 21 (19–24) | 19 (18–21) | < 0.001 |
| Head circumference (cm) | 49.8 (48.8–50.8) | 49.3 (48.4–50.3) | 49.0 (48.0–49.9) | 48.6 (47.8–49.1) | < 0.001 |
| PIQ | 94 (86–105) | 87 (80–101) | 91 (82–101) | 85 (78–91) | 0.007 |
| VIQ | 96 (90–103) | 94 (90–102) | 91 (83–101) | 87 (81–93) | 0.003 |

<sup>1</sup> Values are presented as median (interquartile range), except for age, which is reported as mean (minimum, maximum).
<sup>2</sup> Pearson's Chi-squared test was used for sex distribution; Kruskal–Wallis rank-sum tests were used for continuous variables.

**Fig. 1.**
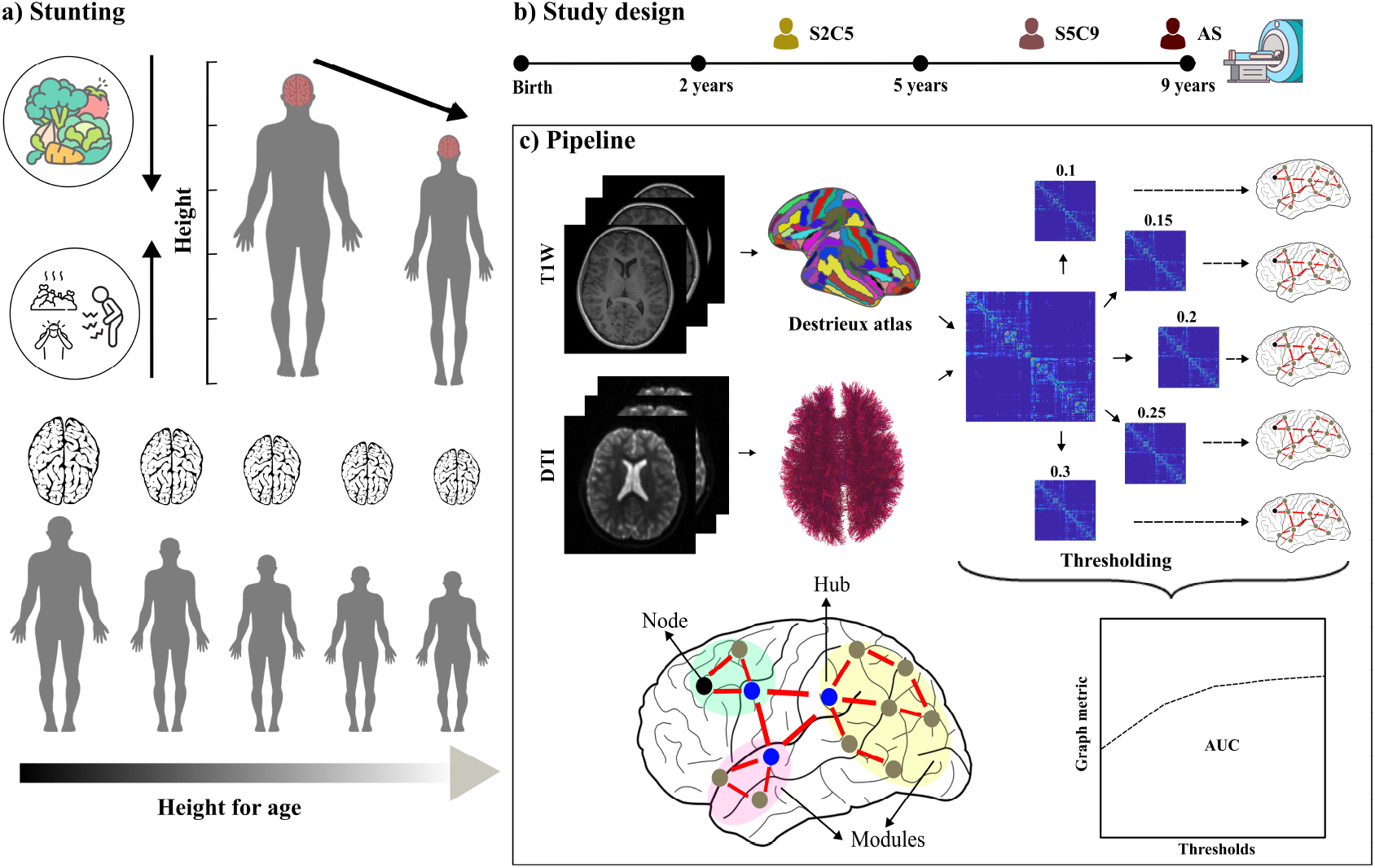
Study overview and structural connectome analysis pipeline. a) Conceptual illustration of stunting and growth faltering across childhood, showing reduced height-for-age and its potential association with altered brain development. The grayscale gradient represents variation in height-for-age across participants. b) Longitudinal study design showing assessment timepoints from birth through early childhood and neuroimaging acquisition at 9 years of age. Early-life measures included socioeconomic and developmental assessments collected at multiple developmental stages. c) Structural connectome processing pipeline. T1-weighted MRI data were used to generate cortical parcellations based on the Destrieux atlas, while diffusion-weighted imaging (DTI) data were used for whole-brain tractography. Structural connectivity matrices were constructed by mapping streamlines between atlas-defined regions. Connectivity matrices were thresholded across a proportional threshold range (10–30%) to evaluate graph theoretical measures across network densities, and area under the curve (AUC) metrics were subsequently derived. Example graph theoretical concepts including nodes, hubs, and modules are illustrated. Abbreviations: S2C5-stunted till two with catch-up at five; S5C9-stunted till five with catch-up at nine; AS-always stunted.

### 2.2 Anthropometric measures

The HAZ was calculated using the Multicentre Growth Reference Study (MGRS) standards, and stunting was defined as HAZ < − 2 SD. Trained, local staff, measured height (in cm) and weight (in kg) at different time points. For all infants until two years of age, an infantometer was used to measure length to the nearest cm and then a stadiometer was used. An electronic weighing scale was used to gauge weight to the nearest 10 grams. All machines were recalibrated periodically, and the local staff underwent regular retraining.

### 2.3 MRI acquisition

All neuroimaging data were acquired while children were awake without anaesthesia in a 3T Siemens Skyra MRI scanner stationed in the Radiology department of CMC Vellore, India. Structural T1-weighted was acquired using the magnetization-prepared rapid gradient-echo (MP-RAGE) sequence with the following parameters: TR = 2350 ms, TE = 1.74 ms, TI = 1100 ms, FOV = 256 and Slice thickness = 1 mm. Children were administered a single dose of Triclofos syrup 0.5ml (50mg)/kg to the maximum of 15 ml (1500mg) if found apprehensive. The project psychologist accompanied and reassured the child during the visit to the radiology department and the MRI scan procedure. If the MRI scan could not be done due to the child’s apprehensions, one more session was attempted within a month. If any child showed apprehensions during both visits, the MRI scan for that child was abandoned.

### 2.4 Preparation of structural connectivity networks

The diffusion MRI (dMRI) data were pre-processed and analysed using the Basic and Advanced Tractography with MRtrix for All Neurophiles (BATMAN) pipeline implemented in the MRtrix3 software package [26]. Pre-processing steps included denoising, eddy current correction, and bias field correction to optimise image quality and reduce imaging artefacts [27, 28]. Fibre orientation distributions were subsequently modelled to estimate white matter structural connectivity between brain regions [29, 30].

Brain parcellation was performed using the FreeSurfer software package based on the Destrieux atlas [31], resulting in 164 cortical, subcortical and whole cerebellar regions of interest (ROIs). Diffusion tractography outputs were mapped onto these ROIs to generate participant-specific structural connectivity matrices, where each matrix element represented streamline-based connectivity strength between pairs of regions (Figure. 1c).

Whole-brain structural connectivity (SC) matrices were constructed for each participant from diffusion MRI tractography, yielding undirected, weighted networks of 164 × 164 region of interests (ROIs). Edge weights reflected streamline-based connectivity strength and were normalised to the range 0–1. Self-connections were removed prior to analysis. All graph-theoretical analyses were performed using the Brain Connectivity Toolbox (BCT) implemented in MATLAB, together with in-house scripts. To minimise bias arising from a single threshold choice, networks were proportionally thresholded across a range of densities (10–30%, in 5% increments), retaining the strongest connections at each density. Graph metrics were computed at each density for subject-level analyses (Figure. 1c).

#### 2.4.1 Definitions of global network metrics

Global network metrics were used to characterise large-scale organisational properties of the structural connectome. These measures capture complementary aspects of network segregation (clustering coefficient, local efficiency, and modularity), network integration (global efficiency and characteristic path length), and network resilience (assortativity and mean nodal strength).

1. **Clustering coefficient (Cp):** The clustering coefficient quantifies the tendency of nodes to form locally interconnected neighbourhoods, reflecting the degree of network segregation and specialised local processing. In weighted networks, the clustering coefficient measures the prevalence of triangular connections around a node, taking into account edge weights, and is averaged across all nodes. For node *i*, the clustering coefficient is defined as 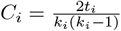, where *t*_*i*_ denotes the weighted number of triangles involving node *i* and *k*_*i*_ denotes its degree. The global clustering coefficient (*C*_*p*_) is obtained by averaging *C*_*i*_ across all nodes. Higher values indicate stronger local clustering and increased network segregation.
2. **Local efficiency (E**_*loc*_): Local efficiency measures the efficiency of information transfer within the immediate neighbourhood of a node and reflects the fault tolerance of the network to local disruptions. It is computed as the global efficiency of each node’s subgraph and averaged across all nodes. Local efficiency is defined as 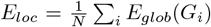, where *G*_*i*_ denotes the subgraph formed by the neighbours of node *i*. Higher values indicate greater robustness and redundancy of local connectivity.
3. **Modularity (Q):** Modularity quantifies the extent to which a network can be subdivided into distinct modules characterised by dense intra-modular and sparse inter-modular connections. For weighted networks, modularity is defined as 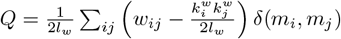, where *w*_*ij*_ represents the edge weight between nodes *i* and *j*, 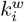 is the weighted degree of node *i, l*_*w*_ is the total weight of the network, and *δ*(*m*_*i*_, *m*_*j*_) equals 1 if nodes belong to the same module. Higher modularity values reflect greater network segregation and specialised functional organisation.
4. **Global efficiency (E**_*glob*_): Global efficiency measures the capacity for parallel information transfer across the entire network and reflects global integration. It is defined as the average inverse shortest path length between all pairs of nodes, 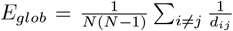, where *d*_ij_ denotes the shortest weighted path length between nodes *i* and *j*. Higher values indicate more efficient long-range communication.
5. **Characteristic path length (L**_*p*_): Characteristic path length represents the average shortest path length between all node pairs and is inversely related to global efficiency. It is defined as 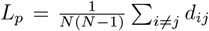. Lower values reflect more efficient global integration, whereas higher values indicate reduced long-range connectivity.
6. **Assortativity:** Assortativity measures the tendency of nodes to connect to other nodes with similar degree or strength. Positive assortativity reflects a preference for high-degree nodes to connect with other high-degree nodes and is considered an indicator of network resilience to targeted attacks. Negative assortativity suggests increased vulnerability of the network.
7. **Mean nodal strength:** Mean nodal strength reflects the average sum of connection weights incident on nodes and provides a measure of overall network connectivity. For node *i*, strength is defined as *s*_*i*_ = Σ _*j*_ *w*_*ij*_, and mean strength is obtained by averaging *s*_*i*_ across all nodes. Higher values indicate stronger overall connectivity and increased network robustness.

#### 2.4.2 Definitions of nodal network metrics

Nodal network metrics were used to characterise region-specific topological properties of the structural connectome. These measures capture the contribution of individual brain regions to network integration, segregation, and inter-modular communication. All nodal metrics were computed on weighted, undirected networks.

1. **Nodal degree:** Nodal degree quantifies the number of connections incident on a node and reflects its level of topological connectedness. For node *i*, degree is defined as *k*_*i*_ = Σ _*j*_ *a*_*ij*_, where *a*_*ij*_ represents the binary adjacency between nodes *i* and *j*. Nodes with higher degree are more densely connected and may play central roles in network communication.
2. **Nodal strength:** Nodal strength is the weighted analogue of degree and represents the sum of connection weights incident on a node. For node *i*, strength is defined as *s*_*i*_ = Σ _*j*_ *w*_*ij*_, where *w*_*ij*_ denotes the weight of the edge connecting nodes *i* and *j*. Higher nodal strength indicates stronger overall connectivity and greater influence on network dynamics.
3. **Betweenness centrality (BC):** Betweenness centrality measures the proportion of shortest paths between all pairs of nodes that pass through a given node, reflecting its importance for global information transfer. For node *i*, betweenness centrality is defined as 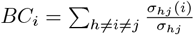, where *σ*_*hj*_ is the total number of shortest paths between nodes *h* and *j*, and *σ*_*hj*_(*i*) is the number of those paths that pass through node *i*. Nodes with high betweenness centrality act as critical bridges within the network.
4. **Participation coefficient (PC):** The participation coefficient quantifies the extent to which a node’s connections are distributed across different network modules. For node *i*, the participation coefficient is defined as 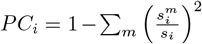, where 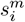 represents the strength of node *i*’s connections within module *m*. Nodes with high participation coefficient facilitate inter-modular communication and are considered connector hubs.
5. **Within-module degree z-score (Z):** The within-module degree z-score measures how well connected a node is relative to other nodes within the same module. It is defined as 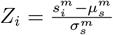, where 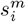 is the within-module strength of node *i*, and 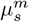 and 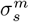 are the mean and standard deviation of strengths within module *m*. Nodes with high z-scores are considered provincial hubs.

#### 2.4.3 Area under the curve (AUC) summarisation

To provide a threshold-independent summary of network topology, all global and nodal graph-theoretical measures were integrated across the predefined density range using AUC analysis. For each participant and each metric, values computed at individual network densities were summarised using numerical integration across the density range. The resulting AUC values were used for all subsequent statistical analyses, reducing sensitivity to single-threshold selection and enhancing robustness.

#### 2.4.4 Hub identification

Network hubs were identified using a multi-criteria nodal ranking approach based on five complementary centrality measures: nodal degree, nodal strength, betweenness centrality, participation coefficient, and within-module degree z-score. All nodal measures were computed at each network density threshold. A node was classified as a hub if it ranked within the top 33% of nodes on at least four out of the five centrality measures, consistent with established criteria for hub identification in structural brain networks [32, 33].

To derive robust group-level hub architecture, hub consistency was assessed across subjects and thresholds. A node was designated as a group-level hub if it was identified as a hub in at least 60% of subjects within that group and was consistently classified as a hub across at least two network density thresholds. This approach ensured that group-level hubs reflected stable and reproducible topological features rather than threshold- or subject-specific effects (Figure 5a).

#### 2.4.5 Statistical analyses

All statistical analyses were conducted using MATLAB (version 2024b) and RStudio. Brain network visualisations were generated using BrainNet Viewer.

##### Edge-wise white matter analyses

To assess group differences in white matter (WM) microstructural and connectivity measures, including fractional anisotropy (FA), mean diffusivity (MD), radial diffusivity (RD), structural connectivity strength (SC), and tract length (TL), edge-wise comparisons were performed between each stunted group (AS, S2C5, S5C9) and the never-stunted (NS) group. For each metric, values corresponding to individual edges were compared using the Wilcoxon rank-sum test. To control for multiple comparisons across edges within each group comparison, false discovery rate (FDR) correction was applied.

##### Distance-based connection analyses

To investigate whether group differences were preferentially expressed across connections of different spatial scales, edges were stratified into short-range (SR), medium-range (MR), and long-range (LR) categories based on the tract length (TL) distribution in the NS group. Edges with TL below the 25th percentile (Q1 = 52.47) were classified as SR, edges above the 75th percentile (Q3 = 110.93) as LR, and edges between Q1 and Q3 as MR (Figure 3a).

For each subject, TL matrices were binarised according to these thresholds to compute the number of SR, MR, and LR connections. To additionally capture connection weights, subject-level SC matrices were multiplied by the corresponding binarised TL matrices, yielding SR, MR, and LR connection strengths. Group differences in both connection counts and strengths were assessed using the Wilcoxon rank-sum test, with FDR correction applied for multiple comparisons.

##### Graph-theoretical analyses

For all global and nodal graph-theoretical measures, area under the curve (AUC) values were computed across the network density range of 10–30%. These AUC values provided a threshold-independent summary metric for each participant and were used for all group-level statistical comparisons.

Group differences in global graph metrics were assessed by comparing AUC values between each stunted group and the NS group using the Wilcoxon rank-sum test. FDR correction was applied separately for each global metric to account for multiple testing.

For nodal graph metrics, AUC values were compared between each stunted group and the NS group on a node-wise basis using the Wilcoxon rank-sum test. To control for multiple comparisons across nodes, FDR correction was applied within each group comparison.

All statistical tests were two-tailed, and statistical significance was defined as *p <* 0.05 after FDR correction.

## 3 Results

The results are organized into four parts. First, between-group differences in white matter (WM) microstructural and tract-based metrics derived from diffusion tensor imaging are reported. Second, alterations in short-, medium-, and long-range structural connections to characterize changes in spatial wiring patterns are examined. Third, group-level differences in global and nodal graph-theoretical properties of structural brain networks are reported, focusing on the notions of segregation, integration, and resilience. Finally, group-level hub architecture are compared to assess whether early childhood stunting and subsequent catch-up growth are associated with reorganization of core networks information processing in the brain.

### 3.1 Between-group differences in metrics quantifying structural connectome across typically developing, stunted and stunted-at-birth children who caught up with normal development trajectory

Group comparisons between never stunted (NS), stunted until two with catch-up at five (S2C5), stunted until five with catch-up at nine (S5C9), and always stunted (AS) children were performed using the Wilcoxon rank-sum test across multiple diffusion tensor imaging (DTI) metrics, including fractional anisotropy (FA), mean diffusivity (MD), radial diffusivity (RD), streamline count (SC), and tract length (TL). Following false discovery rate (FDR) correction for multiple comparisons, no significant group differences were observed for FA, MD, RD, or SC (Figure s1-s4).

In contrast, tract length showed significant between-group differences between NS and AS children that survived FDR correction. The differences indicate longer tract lengths in NS children relative to AS children across these connections (Figure 2). No tract length differences survived correction in comparisons involving the catch-up groups (NS-S2C5 or NS-S5C9).

**Fig. 2.**
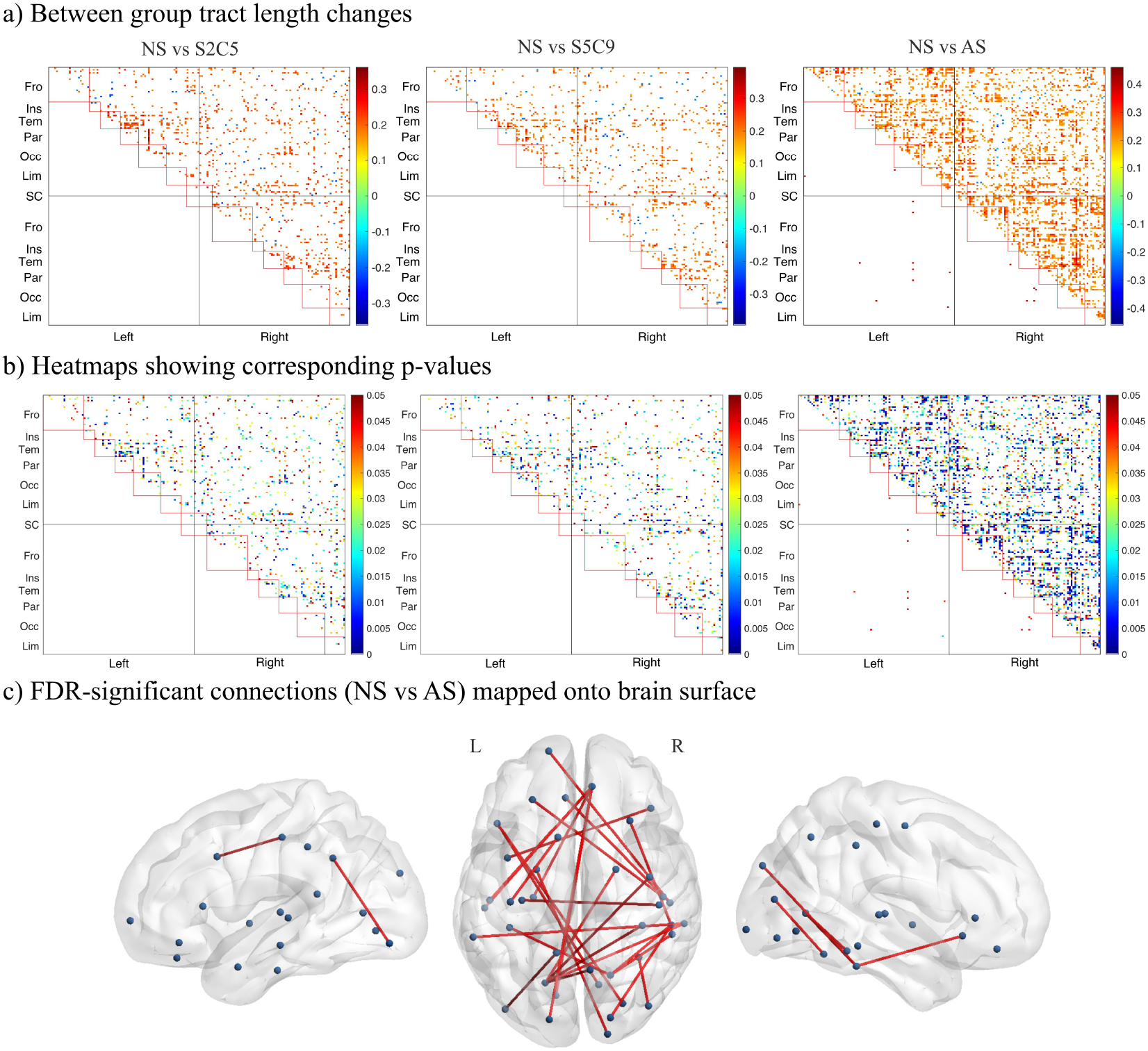
Group differences in tract length (TL). (a) Heatmaps depict groupwise tract length differences for NS vs S2C5, NS vs S5C9, and NS vs AS. Each matrix shows pairwise differences between regions; the upper triangle denotes edges with uncorrected significance (*p <* 0.05), scaled by effect size (Fisher’s 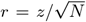). The lower triangle highlights edges surviving FDR correction for multiple comparisons. (b) Heatmaps showing corresponding p-values (c) Connections significantly different between NS and AS after FDR correction, mapped onto the brain surface.

### 3.2 Alterations in short, medium, and long range brain connections

Recent advances in connectomics research indicates short-range and long range functional connectivity have divergent functional outcomes [34]. Hence, to further categorize tract lengths, the following thresholds were used for short-range (SR; < 52.48), medium-range (MR; 52.48–110.94), and long-range (LR; *>* 110.94) connections in NS children (Figure 3a). These thresholds were applied uniformly across all groups. Group comparisons revealed marked differences in the distribution of connections across these spatial ranges (Figure 3b). Compared to NS, AS children exhibited a significant increase in the number of SR (*z* = − 2.64, *p* = .008) and MR (*z* = − 2.68, *p* = .007) connections and a corresponding reduction in LR (*z* = 3.30, *p* =< .001) connections, all of which survived FDR correction. No significant differences were observed for S2C5 children, whereas S5C9 children showed intermediate patterns that did not reach statistical significance for connection counts.

**Fig. 3.**
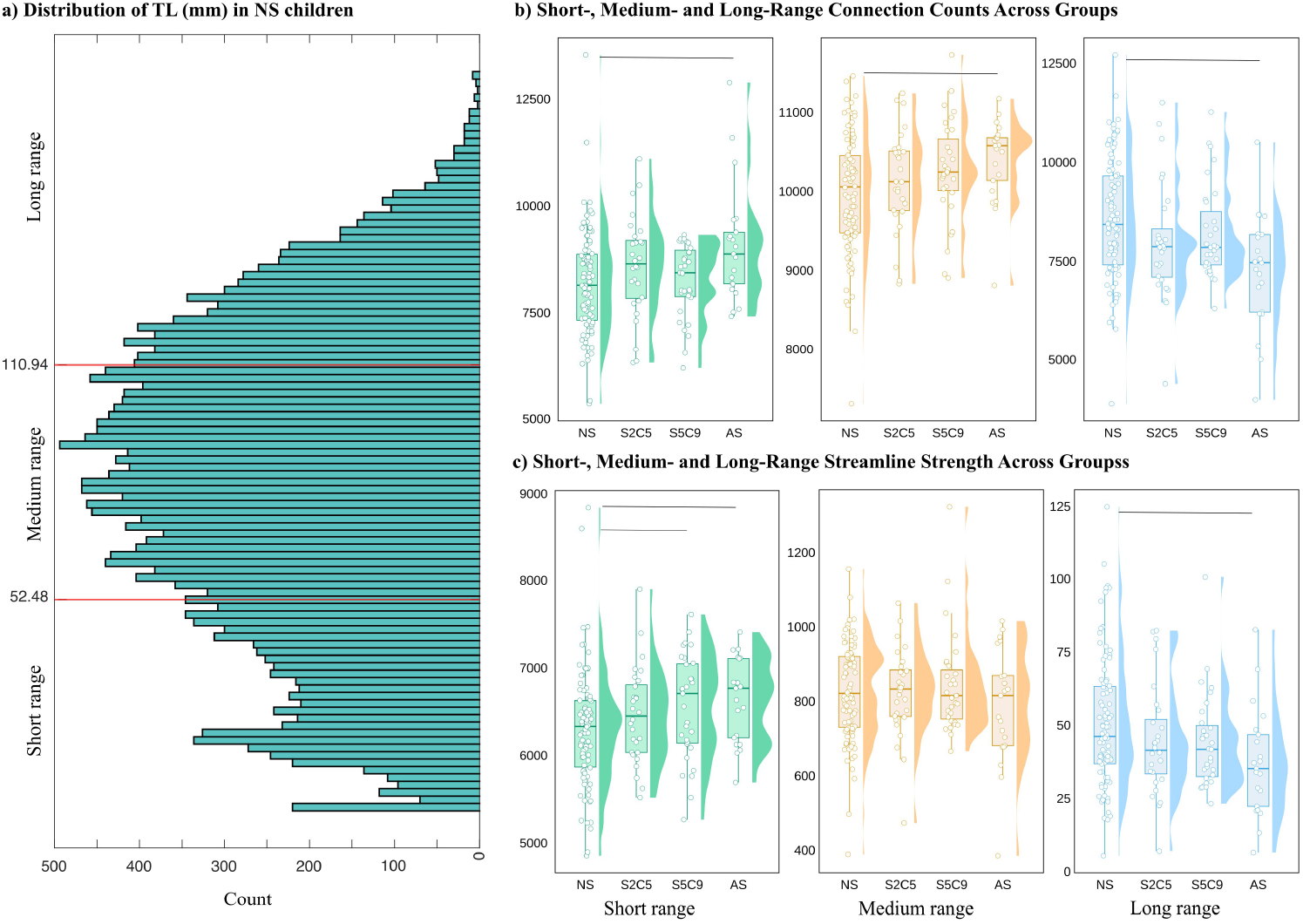
Range-based tract distribution, connection counts, and streamline strength across groups. (a) Histogram of tract lengths in NS children showing thresholds (red lines) defining short-range (SR), medium-range (MR), and long-range (LR) connections. (b) Box and violin plots of the number of SR (green), MR (orange), and LR (blue) connections across groups (NS, S2C5, S5C9, AS). (c) Box and violin plots of total streamline count (SC strength) for SR, MR, and LR connections. Horizontal lines above subplots in panels b and c indicate groups with FDR-significant differences.

Analysis of total streamline count (SC strength) within each range revealed that AS (*z* = − 2.57, *p* = .01) and S5C9 (*z* = − 2.27, *p* = .02) children showed significantly higher SR connection strength relative to NS children (FDR-corrected; Figure 3c). In contrast, LR connection strength was significantly reduced in AS children (*z* = 2.73, *p* = .006) compared with NS children. Together, these findings indicate a shift towards more locally constrained structural connectivity in persistently stunted children, with partial reorganisation evident in children showing later catch-up growth.

### 3.3 Topological organization of brain networks across childhood development spectrum

Across all groups, structural brain networks exhibited small-world organization, densely connected locally with sparse but effective long-range connections that facilitate shorter pathways to reach between physically distant nodes. Mathematically, this is characterized by normalized clustering coefficients greater than 1 (*γ >* 1) and small-worldness indices greater than 1 (*σ >* 1) and higher global and local efficiency (see Methods for details). Although normalised path length (*λ*) was slightly greater than 1 across groups, the overall pattern was consistent with small-world topology (Figure 4a).

**Fig. 4.**
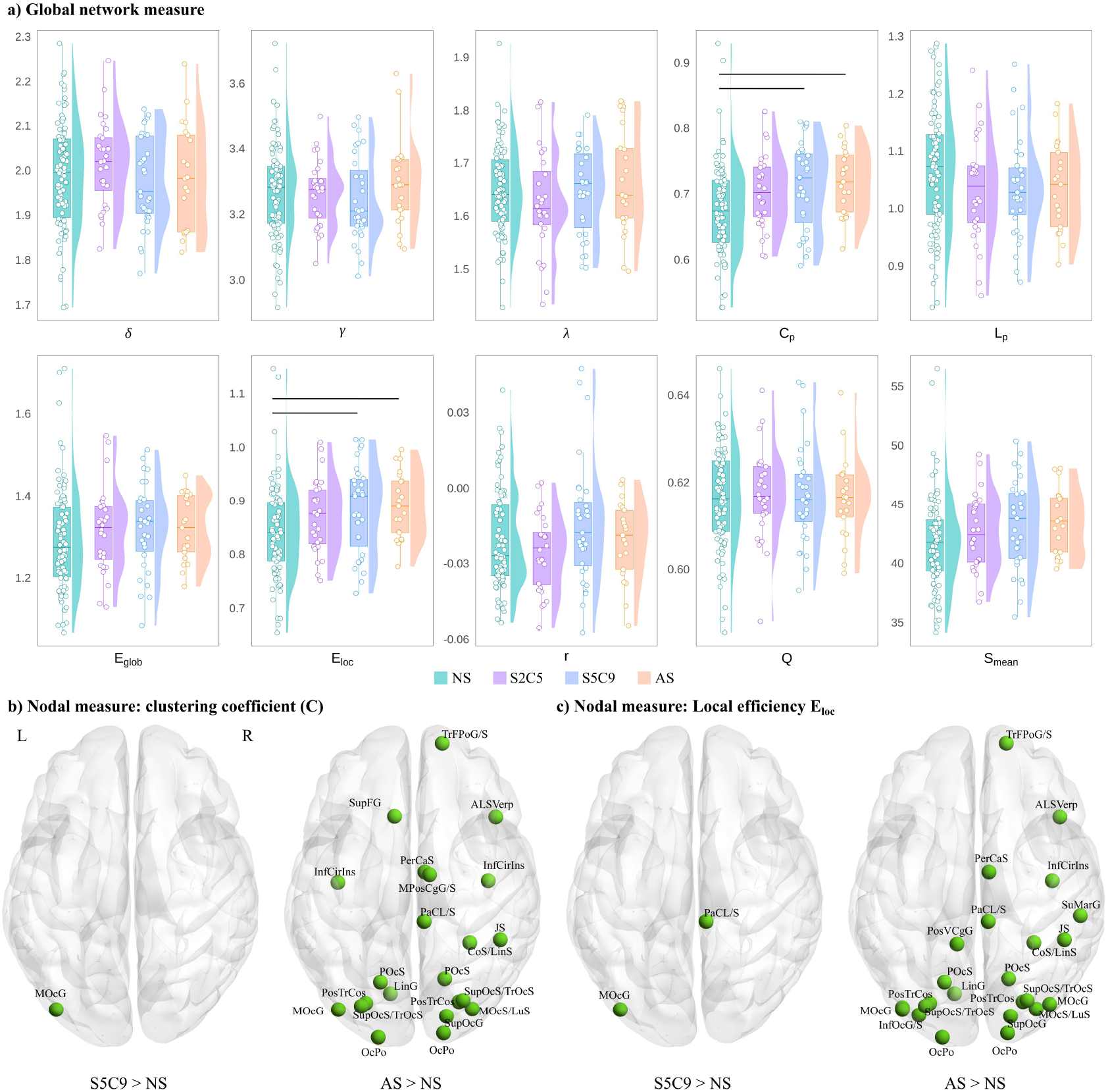
Group differences in brain network topology. (a) Group comparison of global network measures, including small-worldness (*δ*), normalized clustering coefficient (*γ*), normalized characteristic path length (*λ*), clustering coefficient (*C*_*p*_), characteristic path length (*L*_*p*_), global efficiency (*E*_glob_), modularity (*Q*), mean nodal strength (*S*_mean_), and mean local efficiency (*E*_loc_). Boxplots show median, interquartile range, and individual data points. Significant group differences are indicated by horizontal lines. (b) Regional differences in clustering coefficient (*C*) between groups, with significant nodes shown in green. (c) Regional differences in local efficiency (*E*_loc_) between groups, with significant nodes shown in green. L = left hemisphere; R = right hemisphere.

Relative to NS children, both S5C9 and AS groups demonstrated significantly increased clustering coefficient (*C*_*p*_) and mean local efficiency (*E*_*loc*_), indicating enhanced local segregation in the brain networks. Specifically, *C*_*p*_ was higher in S5C9 (*z* = − 2.60, *p* = .009) and AS (*z* = − 2.90, *p* = .004) groups, while *E*_*loc*_ was also elevated in S5C9 (*z* = − 2.48, *p* = .013) and AS (*z* = − 2.69, *p* = .007) groups after FDR correction.

Although mean nodal strength showed nominal increases in S5C9 and AS children relative to NS, these effects did not survive correction for multiple comparisons. No significant group differences were observed for characteristic path length (*L*_*p*_), global efficiency (*E*_*glob*_), assortativity, modularity (*Q*), or mean strength after FDR correction, suggesting preserved global integration and resilience at the whole-network level.

### 3.4 Local regional alterations in network topologies in ECS

At the nodal level, group differences were observed selectively for clustering coefficient and local efficiency (Table s1-s2). Compared with NS children, S5C9 children exhibited increased clustering coefficient in a focal region of the left middle occipital gyrus (*z* = − 3.65, *p* =< .001). In contrast, AS children showed widespread increases in clustering coefficient across 22 nodes spanning frontal, occipital, temporo-occipital, insular, and medial regions (Figure 4b & Table s1).

A similar pattern was observed for local efficiency. S5C9 children showed increased local efficiency in the left middle occipital gyrus and right paracentral lobule/sulcus, whereas AS children exhibited significantly increased local efficiency across 24 nodes distributed across multiple lobes, including occipital, temporo-occipital, insular, parietal, and medial regions (Figure 4c & Table s2).

Overall, nodal alterations in AS children were spatially widespread, reflecting globally increased local segregation, whereas alterations in S5C9 children were more focal and regionally constrained.

### 3.5 Preservation and reorganization of hub architecture

Structural network hubs were identified using five centrality measures: degree, strength, betweenness centrality, participation coefficient, and within-module degree *z*-score. Regions ranking within the top 33% on at least four of the five measures were classified as hubs.

In NS children, hubs were distributed across parietal, occipital, insular, and subcortical regions, including bilateral postcentral sulcus, intraparietal sulcus, superior temporal sulcus, calcarine sulcus, pericallosal sulcus, thalamus, caudate, putamen, hippocampus, and central sulcus (Figure 5b).

**Fig. 5.**
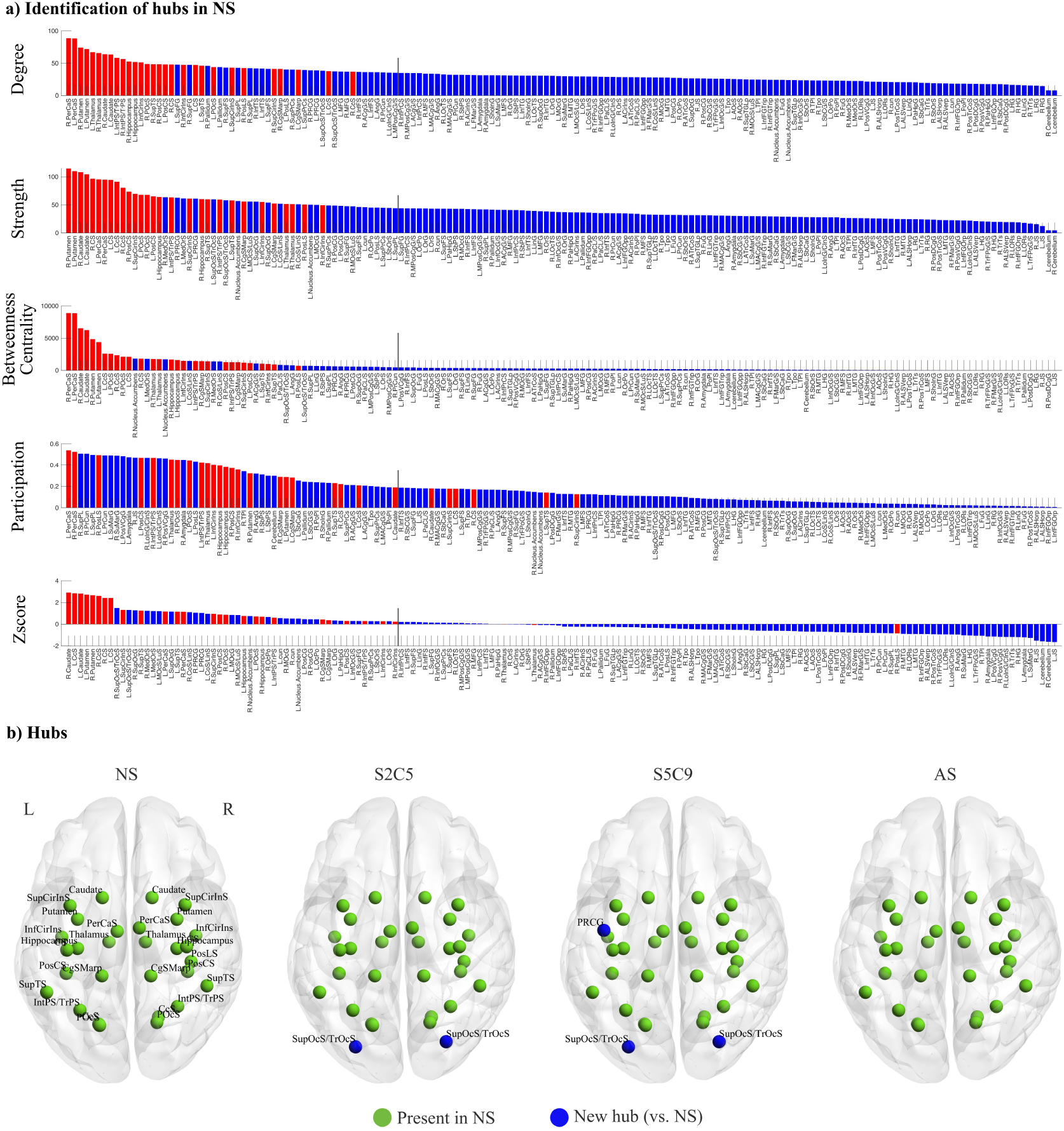
Structural network hubs. a) Node centrality measures (degree, strength, betweenness centrality, participation, and z-score) for the NS group. Bar plots display node scores for each centrality metric, ordered from highest to lowest; the dotted line marks the top 33% of nodes for each measure. b) Brain maps illustrating the spatial distribution of network hubs in each group (NS, S2C5, SSC9, AS). Nodes were classified as hubs if they ranked within the top 33% for at least four out of five centrality measures. Green = hubs present in NS, blue = new hubs (compared to NS).

Hub architecture was largely conserved across all groups. S2C5 children showed all NS hubs, with additional occipital hubs in the bilateral superior occipital sulcus. S5C9 children similarly retained NS hubs but additionally recruited bilateral superior occipital sulcus hubs and a frontal hub in the left precentral gyrus. In contrast, AS children showed no additional hubs beyond those observed in NS children (Figure 5b).

These findings suggest that while persistent stunting is associated with altered local and mesoscale network topology, core hub regions remain preserved. In contrast, catch-up growth, particularly later catch-up is accompanied by recruitment of additional hubs, consistent with compensatory network reorganization.

## 4 Discussion

Overarching goal of this study was to characterize whether early childhood stunting (ECS) and subsequent catch-up growth can be mapped to the organization of cortical white matter measured at late childhood. The primary impact of such understanding will be in development of intervention programs for child education in affected communities and vulnerable groups. Using diffusion MRI and graph-theoretical analyses, the study identifies that early growth disruption is associated with alterations in the spatial and topological organization of white matter connectivity, extending beyond focal tract abnormalities to distributed network-level differences. Together, these findings provide new evidence that ECS is linked to persistent differences in structural brain organization during a critical stage of neurodevelopment, and that catch-up growth may be accompanied by partial, but not necessarily complete—normalization of network architecture. Finally, the differential topological organization of networks based on neurally grounded metrics elucidate the possible mechanisms of compensation of brain function in catch-up growth cohorts, and possible deficiencies that may show up in their behavioral performance based constraints posed by the aberrant network topologies. Thus, grounded in physiological aspects of ECS, understanding developed from this study can be a catalyst for framing educational policies in many regions across the world.

### White matter network organisation as a sensitive marker of early growth disruption

White matter maturation during childhood is characterised by coordinated increases in long-range connectivity, progressive strengthening of integrative hub regions, and refinement of network topology that supports efficient global communication [10, 11, 14]. Such processes can be driven by prolonged axonal growth, myelination, and activity-dependent refinement that extend from infancy through adolescence [10, 11]. In normative development, this reorganisation enables increasingly efficient communication between distributed cortical and subcortical systems and underpins the emergence of higher-order cognitive and socio-emotional functions [35, 36].

Consistent with this developmental framework, our findings indicate that children exposed to ECS exhibit alterations in the balance between local and long-range connectivity, alongside differences in global and nodal network properties. Importantly, such alterations are not readily captured by conventional region-based or tract-specific analyses of MRI, highlighting the value of network-level approaches for understanding how early biological adversity becomes embedded within the developing brain. Long-range connections and hub regions are metabolically costly, develop over extended periods, and depend on tightly regulated myelination and coordinated activity across distributed regions. As a result, these features of network organisation may be particularly sensitive to early growth disruption, leading to persistent differences in integration and efficiency even after physical growth trajectories improve.

At the network level, these maturational changes give rise to small-world topology, characterized by high local clustering indicating a higher *segregation* of functional brain circits combined with short path lengths that support efficient global *integration* [16, 17]. Deviations from this balance, whether through reduced integration measures such as, altered hub centrality, or changes in the spatial distribution of long-range connections, may therefore reflect fundamental disruptions to normative neurodevelopmental processes. Our findings suggest that ECS is associated with such deviations, implicating white matter network organization as a robust marker of ECS and its functional consequences.

Our findings also align with views of the cognition scarcity theory described in resource-constrained settings [37]. Constrained resources, including but not limited to financial scarcity, result in altered cognitive networks enabling quick decisions to meet urgent demands in the environment with subsequent impairments in other networks. This is shown to cause lowering of mental bandwidth limiting cognitive spectrum abilities and tunneling resulting in selective attentional focus and neglect. These network changes can influence economic and livelihood decisions causing perpetuation of poverty [37]. In the same birth-cohort, executive functions such as inhibition and phonemic fluency were shown to be affected in children with persistent stunting with a dose response based on the number of years of stunting [38]. Similar influences in a dose response fashion were noted in verbal and total cognition also in this cohort [2]. It is possible that brain network alterations as shown in this analysis can cause lower overall cognition and higher order executive dysfunction in children with persistent stunting.

### Network-level alterations in the context of prior neuroimaging findings

The present results extend a heterogeneous literature examining the neural correlates of early malnutrition and stunting. Electrophysiological studies, particularly those using EEG and quantitative EEG, have consistently demonstrated long-lasting alterations in neural activity, oscillatory dynamics, and functional connectivity following severe malnutrition in early life, with effects persisting into adulthood [6, 39, 40]. Longitudinal EEG evidence further suggests that faltering growth in infancy may predict later disruptions in functional connectivity and cognitive performance even in the absence of clinically diagnosed malnutrition [41].

Recent morphometric evidence from the current cohort has demonstrated graded reductions in total brain volume, subcortical structures, cerebellar white matter, and posterior corpus callosum thickness across children with normal growth, persistent stunting, and catch-up growth, alongside regionally specific reductions in occipital and subcortical volumes [9]. While these findings indicate that ECS is associated with measurable differences in brain morphology, they do not directly address how such alterations are embedded within large-scale patterns of structural connectivity. The present results extend this work by demonstrating that ECS is also associated with differences in the organization of white matter networks, suggesting that early growth disruption influences not only regional brain structure but also the topology and spatial integration of distributed neural systems.

By capturing network-level properties, graph-theoretical analyses provide a more physiologically interpretable perspective to morphometric approaches, revealing alterations that may persist even when gross anatomical differences appear attenuated following physical recovery. This may help reconcile inconsistencies in prior structural MRI findings, as network organization reflects the coordination of multiple regions and pathways rather than isolated volumetric measures. As such, white matter network topology may represent a particularly robust marker of early developmental disruption, capable of detecting enduring differences in brain organization associated with ECS.

### Catch-up growth and compensatory network reorganization

An important implication of our findings concerns the neurodevelopmental significance of catch-up growth following early stunting. Although catch-up growth is often associated with improved cognitive and behavioural outcomes, emerging evidence suggests that physical recovery does not necessarily entail complete normalization of brain development. In the present study, children who exhibited catch-up growth showed patterns of white matter organisation that were intermediate between persistently stunted and non-stunted peers, consistent with partial recovery at the network level.

One possible interpretation is that catch-up growth is accompanied by a compensatory reorganization of white matter networks via a distinct mechanism rather than a simple return to normative developmental trajectories. Developmental plasticity models propose that when early constraints limit typical maturation, the brain may adopt alternative organizational strategies that support function under altered biological conditions [42, 43]. At the network level, this may manifest as increased reliance on consolidation of local connectivity reflected higher local clustering, redistribution of hub centrality, or preservation of global efficiency through alternative routing strategies. While such reorganization may support adaptive functioning in the short term, it could also confer vulnerability under increasing cognitive, emotional, or environmental demands later in development.

Consistent with this view, prior work has shown that altered hub architecture and deviations in the balance between segregation and integration are common features across neurodevelopmental and psychiatric conditions, even in the absence of gross structural abnormalities [21, 22]. The persistence of network-level differences in children with catch-up growth therefore suggests that early growth disruption may recalibrate the developmental trajectory of white matter organization, with potential long-term implications for mental health and cognitive resilience.

Importantly, the present network-level findings align closely with previously reported cognitive outcomes in this cohort. Prior work demonstrated that children who remained stunted across early and middle childhood exhibited significantly lower verbal and total IQ scores compared to those who were never stunted, whereas children who experienced catch-up growth showed intermediate cognitive performance, outperforming persistently stunted peers [2]. The graded pattern observed in both cognitive outcomes and white matter network organisation suggests a shared underlying developmental mechanism, whereby early growth disruption alters the maturation of large-scale structural connectivity that supports higher-order cognition. In this context, persistent alterations in network integration or hub architecture may constrain information processing efficiency, contributing to enduring cognitive vulnerability, while partial recovery of network organisation in children with catch-up growth may underpin relative cognitive resilience. Together, these converging findings strengthen the interpretation that white matter network topology represents a plausible neural substrate linking early growth trajectories to later cognitive function.

## Limitations and future directions

Several limitations should also be acknowledged for a more nuanced understanding of the key results derived in this study. First, the cross-sectional nature of the neuroimaging data precludes direct inference about developmental trajectories of white matter network maturation. Longitudinal imaging throughout childhood and adolescence will be essential to determine whether the observed network differences reflect delayed maturation, persistent alteration, or compensatory reorganization. Second, diffusion MRI provides indirect estimates of structural connectivity and cannot fully disentangle specific microstructural mechanisms such as axonal density, myelination, or fibre coherence. Future studies integrating advanced diffusion models and myelin mapping techniques that may help pinpointing the rigorous biological substrates underlying network-level differences. Finally, due to time constraints present in a hospital setting, and longer scans were not advisable for a younger cohort - a functional MRI assessment was not possible. Future studies can bridge this gap by collecting resting-state fMRI scans which will allow a direct exploration of the relationship between cognitive markers of ECS with neurally meaningful markers such as the strength of the functional connectivity in different groups - always stunted, never stunted and the catch-up growth cohorts. From a translational perspective, application of AI-ML tools on the diffusion imaging and volumetric analysis within a generalized additive mixed-model framework (GAMM) can be productive to track and predict the cognitive function during early childhood development [44].

In summary, this study demonstrates that ECS is associated with alterations in the spatial and topological organization of white matter structural networks in late childhood. These differences extend beyond focal tract or regional abnormalities and implicate large-scale network architecture as a key neural substrate linking early growth disruption to later neurodevelopment. While catch-up growth may support partial recovery, aspects of network organization appear to remain altered, consistent with compensatory reorganization rather than complete normalization. Together, these findings highlight the value of network neuroscience approaches for advancing mechanistic understanding of early adversity and identifying sensitive markers of neurodevelopmental risk and resilience.

## 5 Acknowledgements

Authors acknowledge the generous support of Computing and Network Facility of NBRC. NBRC Core funds (SB, SS, AB), DBT Dementia Science Programme (AB). The authors thank the children, their families, and the staff of the MAL-ED Network Project. The MAL-ED study was conducted as a collaborative project supported by the Bill and Melinda Gates Foundation, the Foundation for the National Institutes of Health, and the National Institutes of Health/Fogarty International Center (grant no. OPP 47075). The 9-year follow-up of the MAL-ED India cohort was supported by an intermediate clinical and public health research fellowship awarded by the DBT/Wellcome Trust India Alliance to Dr. Beena Koshy (grant no. IA/CPHI/19/1/504611).

