## Supplemental Data 1 for "Neurodevelopmental Inequality arising from Early Childhood Stunting: Evidences from Brain Connectivity"

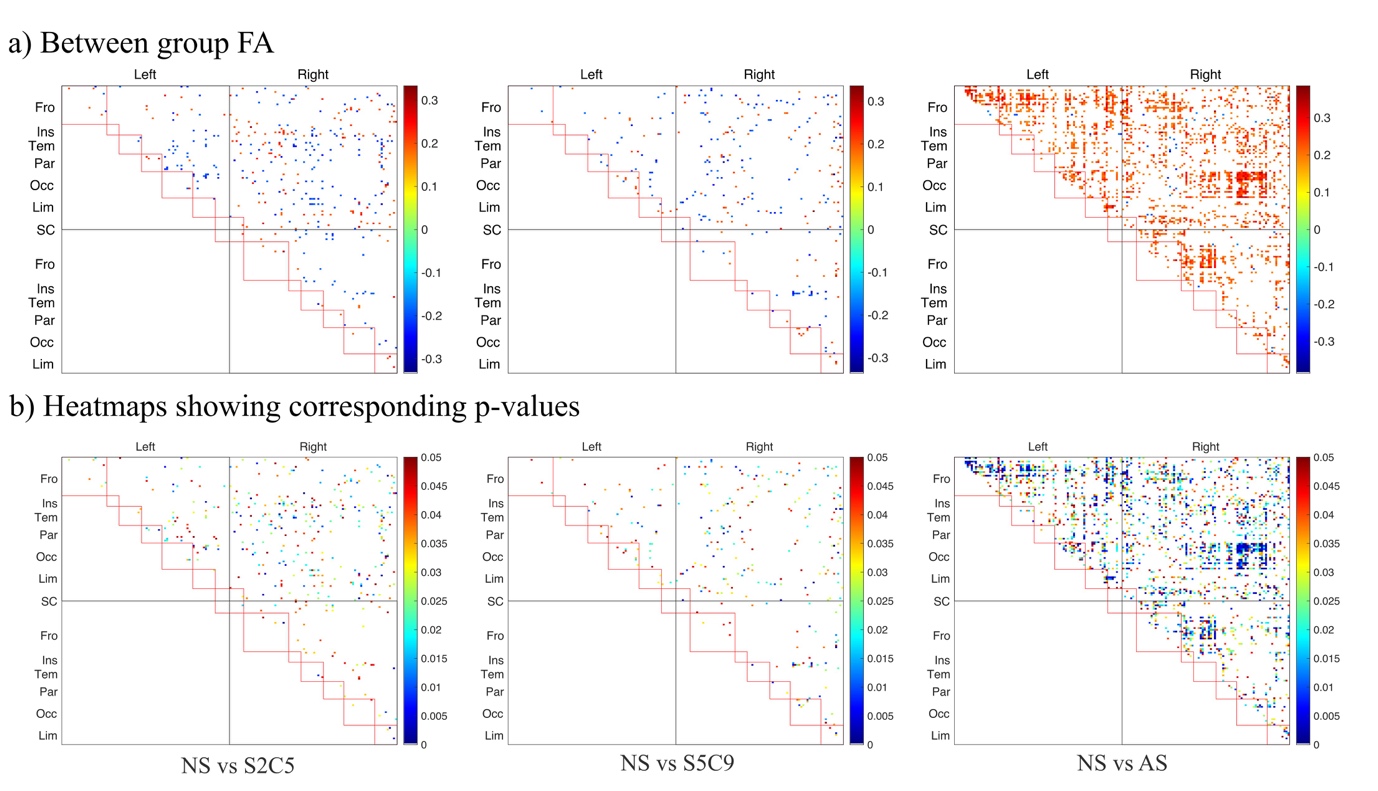

**Figure s1.** Group differences in fractional anisotropy (FA). (a) Heatmaps depict groupwise tract length differences for NS vs S2C5, NS vs S5C9, and NS vs AS. Each matrix shows pairwise differences between regions; the upper triangle denotes edges with uncorrected significance (p < 0.05), scaled by effect size (Fisher’s r = z / √N). The lower triangle highlights edges surviving FDR correction for multiple comparisons. (b) Heatmaps showing corresponding p-values. Abbreviations: NS-never stunted; S2C5- stunted till two with catch-up at five; S5C9- stunted till five with catch-up at nine; AS-always stunted.

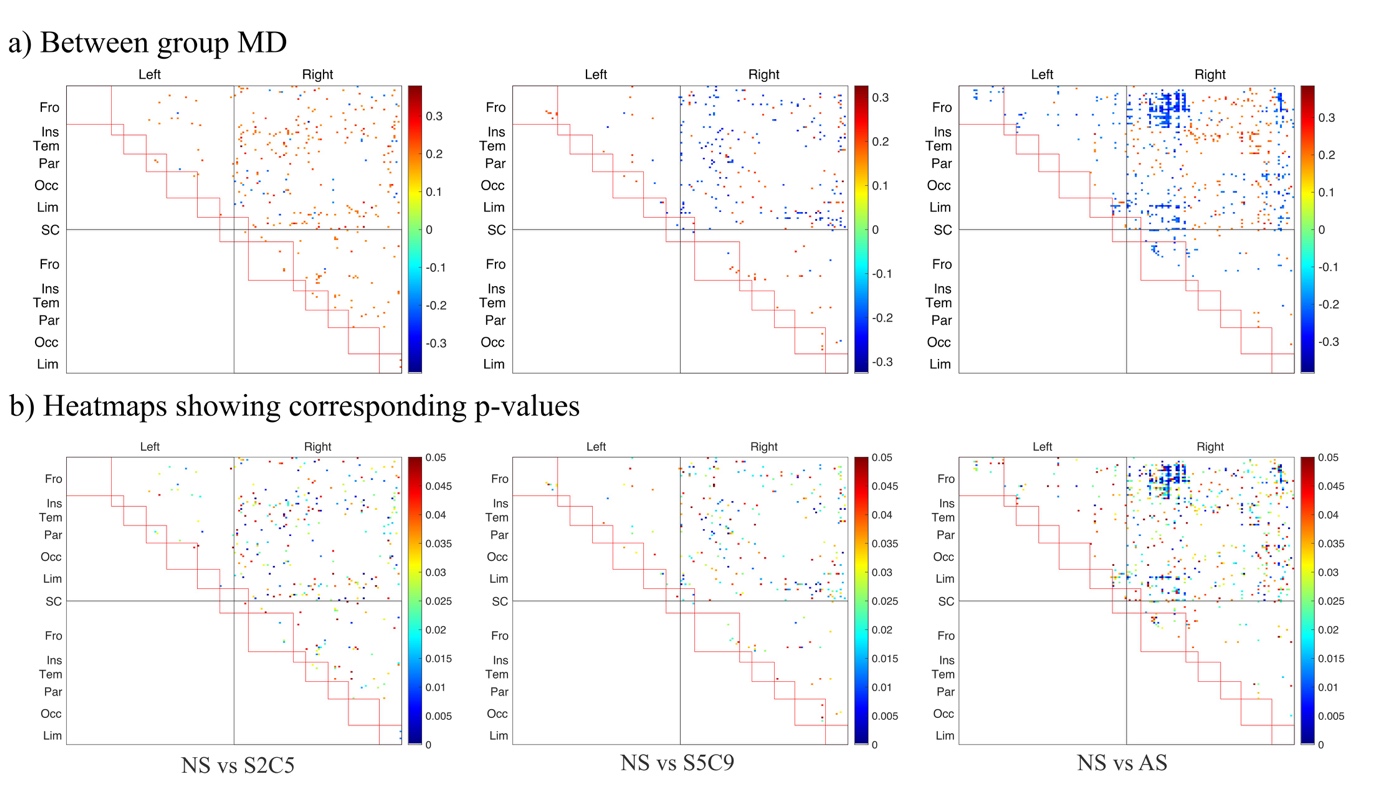

**Figure s2.** Group differences in mean diffusivity (MD). (a) Heatmaps depict groupwise tract length differences for NS vs S2C5, NS vs S5C9, and NS vs AS. Each matrix shows pairwise differences between regions; the upper triangle denotes edges with uncorrected significance (p < 0.05), scaled by effect size (Fisher’s r = z / √N). The lower triangle highlights edges surviving FDR correction for multiple comparisons. (b) Heatmaps showing corresponding p-values. Abbreviations: NS-never stunted; S2C5- stunted till two with catch-up at five; S5C9- stunted till five with catch-up at nine; AS-always stunted.

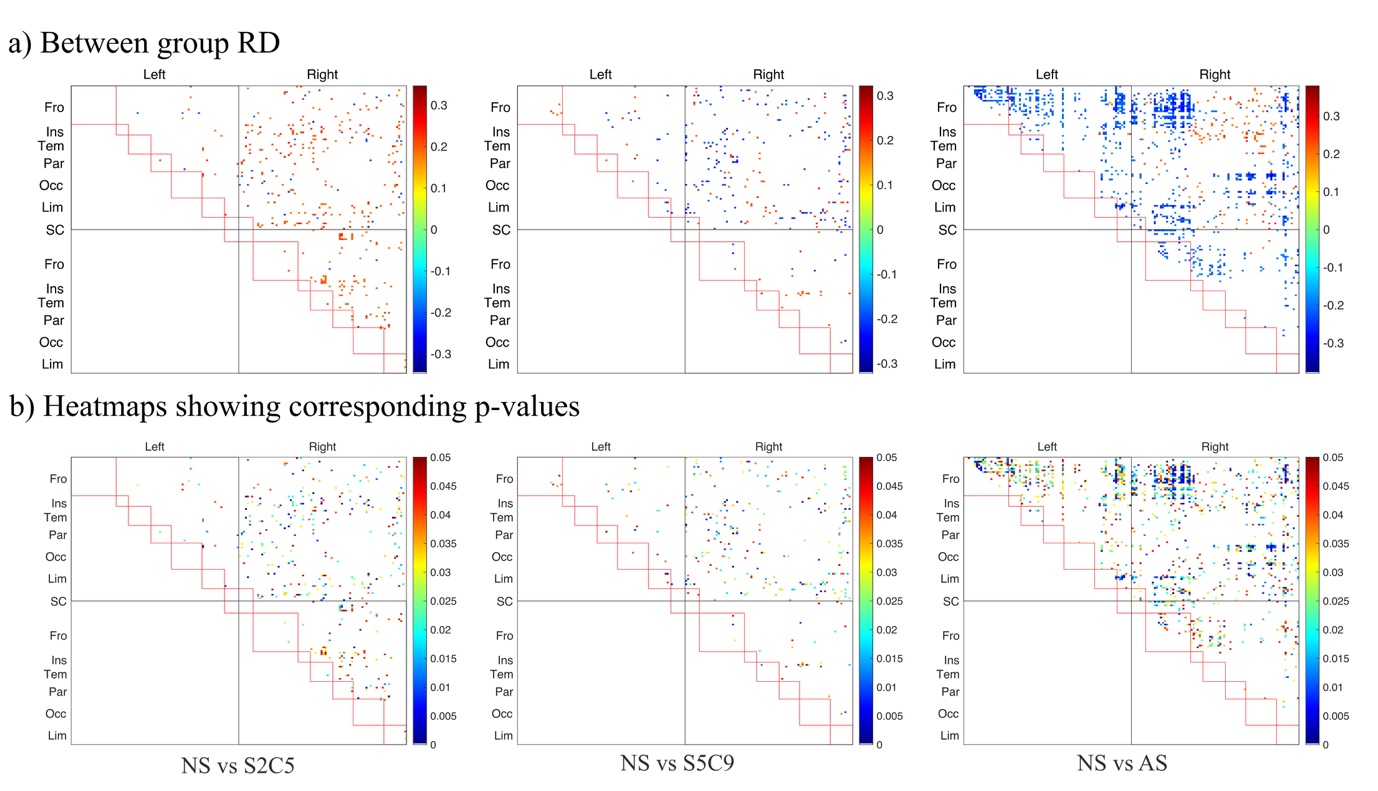

**Figure s3.** Group differences in radial diffusivity (MD). (a) Heatmaps depict groupwise tract length differences for NS vs S2C5, NS vs S5C9, and NS vs AS. Each matrix shows pairwise differences between regions; the upper triangle denotes edges with uncorrected significance (p < 0.05), scaled by effect size (Fisher’s r = z / √N). The lower triangle highlights edges surviving FDR correction for multiple comparisons. (b) Heatmaps showing corresponding p-values. Abbreviations: NS-never stunted; S2C5- stunted till two with catch-up at five; S5C9- stunted till five with catch-up at nine; AS-always stunted.

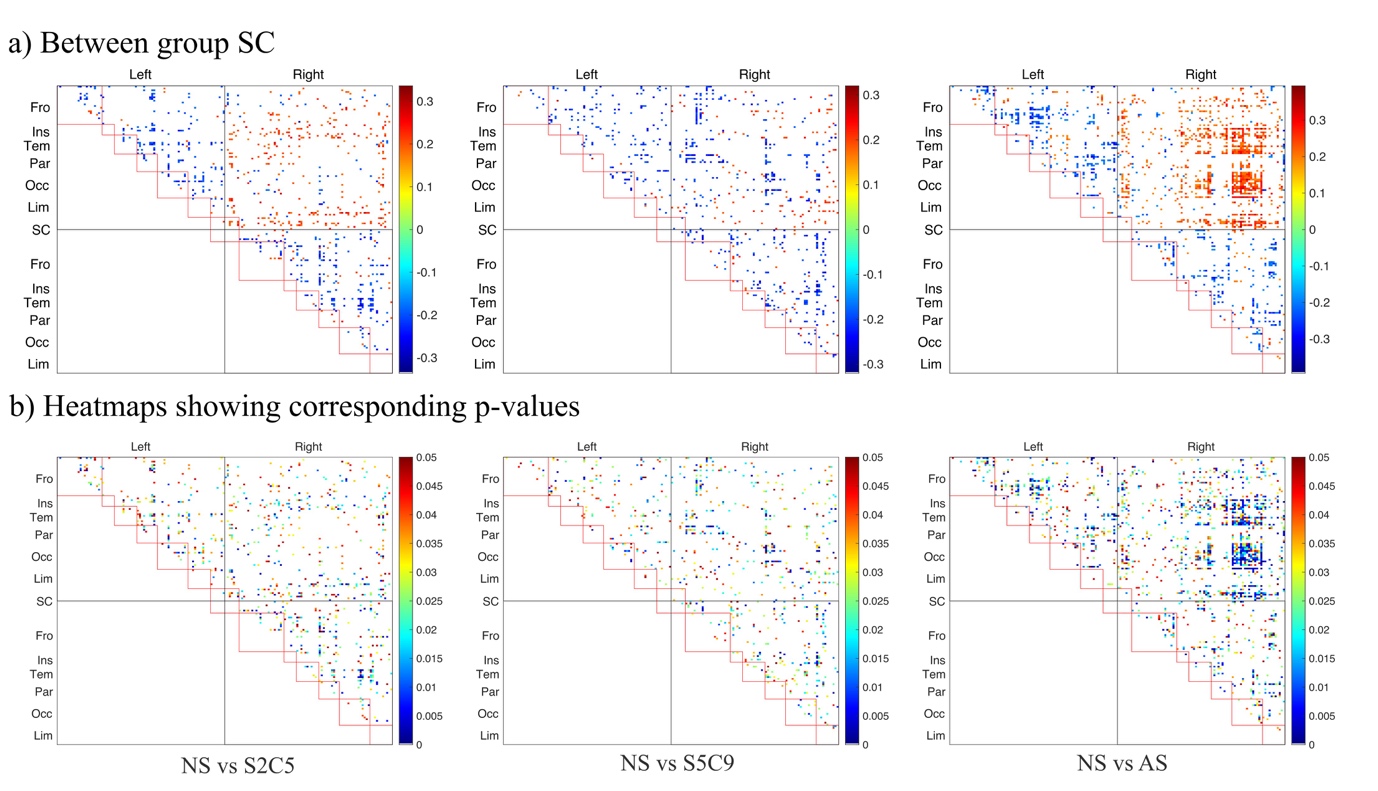

**Figure s4.** Group differences in structural connectivity (SC). (a) Heatmaps depict groupwise tract length differences for NS vs S2C5, NS vs S5C9, and NS vs AS. Each matrix shows pairwise differences between regions; the upper triangle denotes edges with uncorrected significance (p < 0.05), scaled by effect size (Fisher’s r = z / √N). The lower triangle highlights edges surviving FDR correction for multiple comparisons. (b) Heatmaps showing corresponding p-values. Abbreviations: NS-never stunted; S2C5- stunted till two with catch-up at five; S5C9- stunted till five with catch-up at nine; AS-always stunted.

Table s1. Group differences in nodal clustering coefficient

| **ROI** | **NS vs S2C9** | | | **NS vs S5C9** | | | **NS vs AS** | | |
| --- | --- | --- | --- | --- | --- | --- | --- | --- | --- |
|  | **z** | **pval** | **fdr** | **z** | **pval** | **fdr** | **r** | **pval** | **fdr** |
| L.SupFG | -1.20 | 0.231 | 0.511 | -2.35 | 0.019 | 0.134 | -0.26 | 0.006 | **0.047** |
| L.InfCirIns | -2.46 | 0.014 | 0.314 | -2.83 | 0.005 | 0.066 | -0.27 | 0.004 | **0.045** |
| L.SupOcS/TrOcS | -2.44 | 0.015 | 0.314 | -2.76 | 0.006 | 0.066 | -0.26 | 0.006 | **0.045** |
| L.MOcG | -2.19 | 0.028 | 0.319 | -3.65 | 0.000 | **0.042** | -0.31 | 0.001 | **0.038** |
| L.PosTrCoS | -1.64 | 0.102 | 0.368 | -2.84 | 0.004 | 0.066 | -0.33 | 0.001 | **0.038** |
| L.LinG | -0.76 | 0.450 | 0.658 | -2.36 | 0.018 | 0.134 | -0.26 | 0.006 | **0.045** |
| L.POcS | -1.14 | 0.256 | 0.522 | -1.74 | 0.081 | 0.263 | -0.31 | 0.001 | **0.038** |
| L.OcPo | -2.08 | 0.037 | 0.319 | -3.17 | 0.002 | 0.066 | -0.27 | 0.005 | **0.045** |
| R.ALSVerp | 0.58 | 0.563 | 0.737 | 0.16 | 0.875 | 0.891 | -0.26 | 0.006 | **0.045** |
| R.TrFPoG/S | -2.65 | 0.008 | 0.314 | -1.99 | 0.047 | 0.220 | -0.35 | 0.000 | **0.035** |
| R.InfCirIns | -2.07 | 0.039 | 0.319 | -3.05 | 0.002 | 0.066 | -0.32 | 0.001 | **0.038** |
| R.JS | -1.87 | 0.062 | 0.328 | -1.75 | 0.079 | 0.263 | -0.29 | 0.002 | **0.045** |
| R.PaCL/S | -1.99 | 0.047 | 0.319 | -3.28 | 0.001 | 0.066 | -0.29 | 0.003 | **0.045** |
| R.SupOcS/TrOcS | -2.32 | 0.020 | 0.314 | -2.33 | 0.020 | 0.135 | -0.28 | 0.003 | **0.045** |
| R.SupOcG | -1.37 | 0.171 | 0.439 | -1.46 | 0.143 | 0.309 | -0.27 | 0.005 | **0.045** |
| R.MOcS/LuS | -1.12 | 0.262 | 0.522 | -3.03 | 0.002 | 0.066 | -0.27 | 0.004 | **0.045** |
| R.CoS/LinS | -0.71 | 0.477 | 0.680 | -1.73 | 0.083 | 0.263 | -0.26 | 0.006 | **0.045** |
| R.PosTrCoS | -1.52 | 0.130 | 0.422 | -2.75 | 0.006 | 0.066 | -0.27 | 0.004 | **0.045** |
| R.POcS | -2.26 | 0.024 | 0.314 | -2.19 | 0.029 | 0.157 | -0.27 | 0.005 | **0.045** |
| R.OcPo | -1.70 | 0.088 | 0.359 | -2.21 | 0.027 | 0.157 | -0.30 | 0.002 | **0.044** |
| R.PerCaS | -1.76 | 0.078 | 0.333 | -1.65 | 0.098 | 0.279 | -0.27 | 0.005 | **0.045** |
| R.MPosCgG/S | -1.60 | 0.110 | 0.385 | -2.79 | 0.005 | 0.066 | -0.29 | 0.002 | **0.045** |

Table s2. Group differences in nodal local efficiency

| ROI | **NS vs S2C9** | | | **NS vs S5C9** | | | **NS vs AS** | | |
| --- | --- | --- | --- | --- | --- | --- | --- | --- | --- |
|  | **z** | **pval** | **fdr** | **z** | **pval** | **fdr** | **r** | **pval** | **fdr** |
| L.InfOcG/S | -2.17 | 0.030 | 0.310 | -3.24 | 0.001 | 0.050 | -3.01 | 0.003 | **0.033** |
| L.SupOcS/TrOcS | -2.36 | 0.018 | 0.310 | -2.90 | 0.004 | 0.068 | -3.19 | 0.001 | **0.033** |
| L.MOcG | -2.22 | 0.027 | 0.310 | -3.76 | 0.000 | **0.028** | -3.38 | 0.001 | **0.027** |
| L.MOcS/LuS | -3.44 | 0.001 | 0.094 | -2.42 | 0.016 | 0.101 | -3.04 | 0.002 | **0.033** |
| L.PosTrCoS | -1.77 | 0.076 | 0.320 | -2.90 | 0.004 | 0.068 | -3.72 | 0.000 | **0.017** |
| L.LinG | -0.59 | 0.558 | 0.699 | -2.29 | 0.022 | 0.117 | -2.77 | 0.006 | **0.043** |
| L.POcS | -1.36 | 0.175 | 0.390 | -1.96 | 0.050 | 0.165 | -3.60 | 0.000 | **0.017** |
| L.OcPo | -2.07 | 0.039 | 0.310 | -3.19 | 0.001 | 0.050 | -3.05 | 0.002 | **0.033** |
| L.PosVCgG | -1.85 | 0.064 | 0.310 | -1.78 | 0.075 | 0.181 | -2.74 | 0.006 | **0.043** |
| R.ALSVerp | 0.17 | 0.867 | 0.935 | 0.23 | 0.819 | 0.844 | -2.73 | 0.006 | **0.043** |
| R.TrFPoG/S | -2.73 | 0.006 | 0.310 | -1.85 | 0.064 | 0.181 | -3.62 | 0.000 | **0.017** |
| R.InfCirIns | -2.11 | 0.035 | 0.310 | -3.08 | 0.002 | 0.057 | -3.13 | 0.002 | **0.033** |
| R.JS | -1.88 | 0.060 | 0.310 | -1.79 | 0.073 | 0.181 | -3.32 | 0.001 | **0.027** |
| R.SuMarG | -1.26 | 0.207 | 0.409 | -1.23 | 0.218 | 0.337 | -2.89 | 0.004 | **0.043** |
| R.PaCL/S | -2.22 | 0.027 | 0.310 | -3.49 | 0.000 | **0.040** | -3.09 | 0.002 | **0.033** |
| R.SupOcS/TrOcS | -2.04 | 0.041 | 0.310 | -2.43 | 0.015 | 0.101 | -2.74 | 0.006 | **0.043** |
| R.SupOcG | -1.26 | 0.207 | 0.409 | -1.67 | 0.095 | 0.207 | -2.86 | 0.004 | **0.043** |
| R.MOcG | -2.35 | 0.019 | 0.310 | -1.99 | 0.047 | 0.165 | -2.74 | 0.006 | **0.043** |
| R.MOcS/LuS | -1.19 | 0.236 | 0.434 | -2.95 | 0.003 | 0.068 | -3.03 | 0.002 | **0.033** |
| R.CoS/LinS | -0.96 | 0.338 | 0.543 | -1.84 | 0.066 | 0.181 | -2.86 | 0.004 | **0.043** |
| R.PosTrCoS | -1.47 | 0.141 | 0.350 | -2.66 | 0.008 | 0.079 | -2.84 | 0.004 | **0.043** |
| R.POcS | -2.26 | 0.024 | 0.310 | -2.16 | 0.030 | 0.143 | -2.82 | 0.005 | **0.043** |
| R.OcPo | -1.80 | 0.072 | 0.310 | -2.36 | 0.018 | 0.107 | -3.29 | 0.001 | **0.027** |
| R.PerCaS | -2.24 | 0.025 | 0.310 | -2.32 | 0.020 | 0.114 | -2.78 | 0.005 | **0.043** |
